# *Big fish in small tanks:* Stunting in the Deccan Mahseer, *Tor khudree* (Sykes 1849)

**DOI:** 10.1101/2020.04.04.025049

**Authors:** Vishwanath Varma, Rajeev Raghavan, V. V. Binoy

**Author notes:** Correspondence: V.V Binoy, Animal Behaviour and Cognition Programme, School of Natural Sciences and Engineering, National Institute of Advanced Studies (NIAS), Indian Institute of Science (IISc) Campus, Bangalore, India, Email.

## Abstract

The diminished growth and stunting of the Deccan mahseer, *Tor khudree* (Sykes 1849), a mega-fish, endemic to peninsular India is recorded for the first time under high-density laboratory conditions, and its implications for conservation and aquaculture discussed.

---

Mahseers (*Tor* spp.) are freshwater megafish of conservation concern that are facing high levels of population decline (Pinder *et al.*, 2019). As a result, many species of mahseers are being artificially propagated (Keshavanath et al., 2006) in hatcheries, and released into various rivers and reservoirs with an aim of stock enhancement (Ogale, 2002). However, the large size of these fish limits the possibility of laboratory research for improving the outcomes of such restocking interventions. While high density, elevated competition for food resources, and space constraints (Amundsen et al., 2007; Grant & Imre, 2005) have been reported to cause stunting in many fish species, such stunting has not been reported in mahseer.

Deccan mahseer, *Tor khudree* (Sykes 1849) is the most popular and studied mahseer species of the Western Ghats Hotspot, whose local populations have significantly declined in its native range (northern part of the Western Ghats), including being extirpated from their type locality (Goonatilake et al., 2020). However, they have been considered ‘invasive’ in other parts of the Western Ghats (Southern and Central parts), competing with, and resulting in the population decline of other endemic mahseer species (see Pinder et al., 2020, 2015). This fish grows to large sizes in the wild (>1.2 m in Total Length/*T*_L_ and up to 50 kg in Total Weight/*T*_W_, Froese & Pauly, 2019) and is hence of importance to recreational fisheries and aquaculture. These characteristics of deccan mahseer make them a priority for conservation interventions (National Wildlife Action Plan of India, 2019). Here, we report a serendipitous observation that *Tor khudree* maintained under standard laboratory conditions exhibit stunted growth.

Fingerlings of *T. khudree* (4.18 ± 0.08 cm Standard Length/*S*_L_ and 60 day old) were obtained from the hatchery of the Department of Fisheries, Government of Karnataka at Harangi, Kodagu, India. In the laboratory, they were divided into groups of eight individuals and maintained in aquaria of dimensions 40 × 25 × 27 cm for observing behavioural patterns, which is a major knowledge gap in the mahseer literature. Water level was maintained at a height of 25 cm and steel grills covered with mosquito nets were placed as lids to prevent fish from jumping out. Constant aeration was provided to avoid reduction of dissolved oxygen in the tank. There was no restriction on the provision of food, the fish were fed every evening with Taiyo food pellets *(ad libitum),* and water was changed every ten days. Twenty-five tanks were prepared following this protocol, and maintained at room temperature under a natural photoperiod of 12L:12D cycle for a period of 12 months. At the end of the experiment, 50 randomly selected individuals were measured for length (*S*_L_) and weight (*T*_W_).

Fish maintained under laboratory conditions for 12 months exhibited stunting and grew to an average size of 5.52 ± 0.13 cm *S*_L_ (mean ± SE) and 4.08 ± 0.24 g *T_W_,* in contrast to the studies of Singh et al. (2012) who reported an increase of 1 g in body weight (in individuals with average weight 0.10 ± 0.12 g), when reared on high-protein diets over 60 days and Sangma & Basavaraj (2012) who found an average increase of 9.72 cm *T*_L_ and 88.8 g in 120 days in concrete tanks (500 × 500 × 100 cm). A specific growth rate of 0.03 mm/day was observed in our study (Table 1) against 0.81 mm/day by Sangma & Basavaraj (2012). In natural waters, *T. khudree* is known to grow to lengths of 38.3–120.2 mm *T*_L_ in a year (Raghavan et al., 2011).

**Table 1.** Growth, survival and condition factor of the Deccan mahseer, *Tor khudree*, reared in the laboratory condition for 12 months

| Parameters | Value |
| --- | --- |
| Initial length (cm) | 4.18 |
| Final length (cm) | 5.52 |
| Specific Growth Rate (mmday<sup>-1</sup>) | 0.03 |
| Survival (%) | 97 |
| Condition Factor (*K*) | 2.35 |

Fulton’s condition factor *K* (Froese, 2006) of the stunted individuals was high (2.35 in our study *vs.* 1.68, estimated from Sangma & Basavaraja, 2012), indicating that high density rearing does not negatively impact the body condition of *T. khudree* (Table 1). Similarly, the survival rate of *T. khudree* reared at high density in the laboratory was 97%, which was not different from the mortality reported at lower densities under captive conditions (Sangma & Basavaraj, 2012).

Stunting is primarily thought to be caused by socio-environmental constraints experienced by the fish, with little role of genetics (Mohapatra et al. 2017). Nevertheless, such retarded growth has not been described in mahseer populations in either natural or captive environments. The diminished growth and stunting exhibited by *T. khudree* in laboratory conditions may be the result of high-density rearing conditions (8 individuals/tank), since density has been observed to be a husbandry-related stressor in other cyprinids (Nandeesha *et al.,* 2013). Though lack of exercise, which is essential for increasing both food intake and food conversion (Jobling et al., 1993), may be considered a potential cause for stunting, preliminary studies that quantified activity levels of *T. khudree* in laboratory conditions revealed high activity throughout the day (Varma et al., Unpubl.).

Stunting of *T. khudree* could have wider implications for both its farming and conservation. For example, farmers in Andhra Pradesh, India, stunt fingerlings of Indian carps for six to twelve months by subjecting them to high rearing densities (up to 100,000/ha and feeding at a rate of 2–3% of body weight; Nandeesha et al., 2013). Stunted fingerlings exhibit greater feeding, feed conversion efficiency and faster weight gain when they are subsequently transferred to lower densities and improved environmental conditions in production ponds (Nandeesha et al., 2013). If *T. khudree* is capable of such compensatory growth similar to other cyprinids, stocking large numbers of fingerlings in a cost-effective manner and making them available throughout the year for farmers could help in promoting the culture of this species in artificial and natural waters. Additionally, such stunted individuals may be more suitable for ranching and restocking programs due to the ease of transportation from hatcheries to natural habitats, and potential increase in growth rate after release. Miniaturized *T. khudree* could also evolve into a popular aquarium fish due to their morphological features (e.g. bright and large scales), coloration during their juvenile phases, highly active nature under captive conditions, and symbolic status in many states of India (Dinesh et al., 2010).

Providing ‘life skills’ training such as familiarizing with natural prey items and fine-tuning predator avoidance behaviour may be required to prevent early mortality of hatchery reared individuals that are released into the wild (Sloychuk et al., 2016). Stunted *T. khudree* are suitable for experimental arenas and holding conditions that are used for testing popular fish model systems in the laboratory (such as guppy, medaka, zebrafish etc.). Further studies on the morphology, behaviour, physiology and fitness traits of stunted mahseers could improve the outcomes of conservation efforts and restocking interventions.

## ACKNOWLEDGEMENTS

The authors are grateful to the Karnataka State Fisheries Department for kindly providing the fingerlings, and to Darshan Pramod (District Fisheries Department, Kodagu), H. Vasoya, A. Jain, J. Vijayan and A. Singh for their support during the study. VV thanks the Department of Biotechnology, Government of India, for a Research Associate Fellowship. The care and use of experimental animals complied with Indian animal welfare laws, guidelines and policies as approved by the National Institute of Advanced Studies (NIAS) Research Ethics Committee (NIAS/EC/VV/2019). The authors declare no conflict of interest.

## DATA AVAILABILITY STATEMENT

The data that support the findings of this study are available from the corresponding author upon reasonable request.

## REFERENCES

Amundsen, P. A., Knudsen, R., & Klemetsen, A. (2007). Intraspecific competition and density dependence of food consumption and growth in Arctic charr. Journal of Animal Ecology, 76, 149–158. doi:10.1111/j.1365-2656.2006.01179.x

Dinesh, K., Nandeesha, M. C., Nautiyal, P., & Aiyappa, P. (2010). Mahseers in India: A review with focus on conservation and management. Indian Journal of Animal Sciences, 80, 26–38.

Froese, R. (2006). Cube law, condition factor and weight–length relationships: history, meta analysis and recommendations. Journal of Applied Ichthyology, 22, 241–253. doi:10.1111/j.1439-0426.2006.00805.x

Froese, R. and Pauly, D. (2019, February 28). FishBase. World Wide Web electronic publication. Available at www.fishbase.org.

Grant, J. W. A., & Imre, I. (2005). Patterns of density dependent growth in juvenile stream? dwelling salmonids. Journal of Fish Biology, 67, 100–110. doi:10.1111/j.0022-1112.2005.00916.x

de Alwis Goonatilake, S., Fernado, M. & Kotagama, O. (2020, April 7). Tor khudree. The IUCN Red List of Threatened Species Version 2020. Available at https://dx.doi.org/10.2305/IUCN.UK.2020-1.RLTS.T169609A60597571.en.

Jobling, M., Baardvik, B. M., Christiansen, J. S., & Jørgensen, E. H. (1993). The effects of prolonged exercise training on growth performance and production parameters in fish. Aquaculture International, 1, 95–111. doi:10.1007/BF00692614

Keshavanath, P., Gangadhara, B., Basavaraja, N., & Nandeesha, M. C. (2006). Artificial induction of ovulation in pond-raised mahseer, Tor khudree using carp pituitary and Ovaprim. Asian Fisheries Science, 19, 411–411.

Ministry of Environment, Forests and Climate Change. (2017). India’s National Wildlife Action Plan. Available at http://www.indiaenvironmentportal.org.in/files/file/nwap_2017_31.pdf.

Mohapatra, S., Chakraborty, T., Reza, M. A. N., Shimizu, S., Matsubara, T., & Ohta, K. (2017). Short-term starvation and realimentation helps stave off Edwardsiella tarda infection in red sea bream (Pagrus major). Comparative Biochemistry and Physiology Part B: Biochemistry and Molecular Biology, 206, 42–42.

Nandeesha, M. C., Sentilkumar, V., & Antony Jesu Prabhu, P. (2013). Feed management of major carps in India, with special reference to practices adopted in Tamil Nadu. In On-farm Feeding and Feed Management in Aquaculture (Hasan, M. R. & New, M. B. eds) pp. 433–433. FAO Fisheries and Aquaculture Technical Paper, Rome, Italy: FAO.

Ogale, S. N. (2002). Mahseer breeding and conservation and possibilities of commercial culture. The Indian experience. In Coldwater Fisheries in the Trans Himalayan Countries (Petr, T. & Swar, D. B. eds) pp. 193–212. FAO Fisheries Technical Paper, Rome, Italy: FAO.

Pinder, A. C., Britton, J. R., Harrison, A. J., Nautiyal, P., Bower, S. D., Cooke, S. J., … & Walton, S. (2019). Mahseer (Tor spp.) fishes of the world: status, challenges and opportunities for conservation. Reviews in Fish Biology and Fisheries, 29, 417–417.

Pinder, A. C., Raghavan, R., & Britton, J. R. (2015). The legendary hump-backed mahseer Tor sp. of India’s River Cauvery: an endemic fish swimming towards extinction? Endangered Species Research, 28, 11–11.

Pinder, A. C., Raghavan, R., & Britton, J. R. (2020). From scientific obscurity to conservation priority: Research on angler catch rates is the catalyst for saving the hump backed mahseer Tor remadevii from extinction. Aquatic Conservation: Marine and Freshwater Ecosystems, 30, 1809–1809.

Raghavan, R., Ali, A., Dahanukar, N., & Rosser, A. (2011). Is the Deccan Mahseer, Tor khudree (Sykes, 1839) (Pisces: Cyprinidae) fishery in the Western Ghats Hotspot sustainable? A participatory approach to stock assessment. Fisheries Research, 110, 29–29. doi:10.1016/j.fishres.2011.03.008

Sangma, K. O. N., & Basavaraja, N. (2012). Comparative growth and survival of hatchery produced and wild fingerlings of Deccan mahseer, Tor khudree (Sykes). Indian Journal of Fisheries, 59, 89–93.

Singh, S. K., Mishra, U., & Roy, S. D. (2012). Effect of Feeding Enriched Formulated Diet and Live Feed on Growth, Survival and Fatty Acid Profile of Deccan Mahseer, Tor Khudree (Sykes) First Feeding Fry. Journal of Aquaculture & Research Development, 3(6), 1000143. doi:10.4172/2155-9546.1000143

Sloychuk, J. R., Chivers, D. P., & Ferrari, M. C. (2016). Juvenile lake sturgeon go to school: Life ? skills training for hatchery fish. Transactions of the American Fisheries Society, 145, 287–294. doi:10.1080/00028487.2015.1123183

